# Targeting pioneer transcription factor Ascl1 to promote optic nerve regeneration

**DOI:** 10.1101/2023.07.20.549959

**Authors:** Bryan C Dong, Ximei Luo, Cheng Qi, Jiang Qian, Cheng Qian, Feng-Quan Zhou

## Abstract

In adult mammalian central nervous system (CNS) neurons, axon regeneration after injury remains limited due to unfavorable gene regulatory programs. Factors enabling comprehensive epigenetic and transcriptional transitions, for instance, pivotal transcription factors that mediate neurogenesis and morphogenesis may be sufficient to promote CNS axon regeneration. Based on the analyses of multiple public whole-genome RNA and chromatin accessibility sequencing dataset of mouse retina development, as well as previous functional studies on the regeneration-capable dorsal root ganglion neurons, we hypothesize that the overexpression of pioneer transcription factor Achaete-Scute homolog 1 (Ascl1) would promote axon regeneration in the adult mammalian CNS neurons. We employed the optic nerve crush in mice, a common model for studying CNS axon regeneration, neuron survival and glaucoma, to investigate the effect of Ascl1 overexpression on the post-injury optic nerve regeneration. We found that Ascl1 could sufficiently promote regenerated axons past the crush site and significantly preserve the survival of retinal ganglion cells. Mechanistically, we revealed that effects of Ascl1 was mediated by known pro-regeneration factor Sox11 but not others. Together, our study established an effective workflow combined with the integrated computational inference and experimental validation for discovering functionally important target for promoting CNS neuron axon regeneration and survival.

## INTRODUCTION

Axon regeneration of mature neurons in the mammalian central nervous system (CNS) remains virtually absent, presenting a significant biomedical challenge in remediating CNS neural injuries and various neurodegenerative disorders (Qian and Zhou, 2020; Yang et al., 2020). Long-distance axon regeneration is limited by both the inhibitive environment and the lack of intrinsic capacities. Physical barriers, such as post-injury glial scars and debris remnants (Cregg et al., 2014; Neumann et al., 2008), as well as inhibitory cues, such as some myelin associated factors (Giger et al., 2010; Yiu and He, 2006), all constrict axonal extension past the injury site. However, recent studies have indicated that the removal of scarring tissue in either the spinal cord or cerebral cortex (Anderson et al., 2016; Canty et al., 2013), was not sufficient to promote long-distance axon regeneration in CNS. The prevailing consensus is that global gene network regulations, which enable transitions in both transcription and protein translation toward the regeneration state (Qian and Zhou, 2020; (Curcio and Bradke, 2018; Mahar and Cavalli, 2018; Varadarajan et al., 2022; Weng et al., 2017; Williams et al., 2020)), will be necessary to coordinate the long-distance axon regeneration in mature CNS.

One strategy to predict new factors that can promote axon regeneration depends on analyzing the hub regulators involved in the CNS neurons development and maturation, which represent a period requiring comprehensive innate programs for robust axon projection. Until the establishment of synaptic connections, the developed CNS circuitry greatly attenuates redundant neurite extension and branching, which is an evolutionary feature of the mammalians especially (Yang et al., 2022). Along this theory, several transcription factors with obvious up- or down-regulation during neural development have been identified as effective regulators of adult axon regeneration. For example, Sox11 in the Sry-related HMG box family and Klf6 and 7 in the Krüppel-like family have been detected down-regulated across the embryonic-to-postnatal stages. Being consistent with the theory, viral overexpression of Sox11 or Klf6(7) in adult neurons augments post-injury axon regeneration in the optic nerve or the corticospinal tract (CST) (Hilton and Bradke, 2017; Moore et al., 2009; Wang et al., 2015). On the other hand, deletion of Klf4 or Klf9, which is strikingly upregulated along neural maturation, successfully promoted post-injury axon regeneration (Moore et al., 2009).

Similar to the developmental axonal growth, axon regeneration is an orchestra of multiple cellular processes including but not limited to bioenergetics, lipids and protein biosynthesis, cytoskeleton construction, as well as axonal transports (Blanquie and Bradke, 2018; Huebner and Strittmatter, 2009; Wang et al., 2020). These functional executants are termed terminal regeneration associated genes (RAGs) (Huebner and Strittmatter, 2009; van Kesteren et al., 2011), comparing to the upper hierarchal hubs which regulate multiple RAGs. Epigenetic regulators likely play an important role in controlling multiple RAGs loci by globally regulating the chromatin accessibility (Lilja et al., 2013; van Steensel, 2011). Indeed, experimental evidence has been showing that inhibiting histone deacetylase (HDAC) enhances axon regeneration (Cho et al., 2013; Finelli et al., 2013), while EP300 a histone acetyltransferase (HAT) supports the spontaneous axon regeneration of peripheral nervous system (PNS) (Gaub et al., 2011; Lee and Workman, 2007). In addition, recent evident has shown that enhancer of zeste homolog 2 (Ezh2) methyltransferase both facilitates PNS axon regeneration and sufficiently promotes CNS post-injury axon regeneration (Wang et al., 2022).

Logically, there should be a machinery that navigates global epigenetic modifiers to specific motif-dependent gene loci. The pioneer transcription factor (pTF) is a distinct class of transcription factor that can insert into high nucleosome occupancy sites of the closed heterochromatin, and recruiting complex with histone modifiers to initiate epigenetic landscape remodeling (Mayran and Drouin, 2018; Zaret and Carroll, 2011). Among identified pTFs, the basic helix-loop-helix (bHLH) family member Achaete-scute homolog 1 (ASCL1) has been shown to direct vertebrate pro-neuronal differentiation (Raposo et al., 2015; (Liu et al., 2018)), for example, in the spinal cord (Di Bella et al., 2019), and within the cerebral cortex and basal ganglia (Casarosa et al., 1999; Castro et al., 2011). In addition to the function in developmental neurogenesis, Ascl1 has been shown to be one essential factor involved in glia-to-neuron trans-differentiation (Jorstad et al., 2017; Pollak et al., 2013) or fibroblast-to-neuron conversion (Luo et al., 2018), suggesting its ability in remodeling the epigenome of cells to achieve transitions of cell identifies and states. Supported by another evidence, Ascl1 has been shown to act as the key regulator that controls the enhanced spontaneous nerve regeneration in the CAST/Ei mouse strain than in C57BL/6 WT mice. In line with this, Ascl1 overexpression in dorsal root ganglion (DRG) neurons promoted neurite outgrowth ability (Lisi et al., 2017). Together, all evidence suggests the role of Ascl1 in a comprehensive regulation of the genome network enables its function in the pro-neuronal neurogenesis and trans-differentiation, as well as in the post-injury nerve regeneration in mature neurons.

The function of Ascl1 in restoring axon regeneration in the mature CNS neurons remains unknown. In this study, by re-analyzing publicly available RNA- and the chromatin accessible sequencing datasets across different developmental and maturation stages of mouse retina tissue, we identified that at transcripts level many known axon regeneration enhancing genes gradually decrease during retina maturation, including but not restricted to Sox11, Klf6(7), Lin28 and Ezh2. We then applied advanced integrated bioinformatics on multi-omics data and identified many transcription factors as potential inducers for CNS axon regeneration. Based on a newly established scoring method using TF-footprints on the promoter or enhancer regions of a set of known regeneration-associated genes, the novel target Ascl1 was highly ranked. In the functional validation, overexpression of Ascl1 in mouse retina significantly enhanced optic nerve regeneration and RGCs survival after the optic nerve crush. Moreover, we revealed that Sox11 as a potential downstream mediator of Ascl1. Collectively, by combining advanced bioinformatics analysis and optic nerve regeneration model our study demonstrated the role of Ascl1 in promoting CNS neuron survival and axon regeneration.

## RESULTS

### Computational analysis on mRNA and chromatin accessibility sequencing datasets of mouse retina development predicts Ascl1 as a hub directing axon regeneration

Developmental axon elongation of the retinal ganglion cells (RGCs), through being bundled in the optic nerves to reach the brain targets, can achieve a maximal rate at the postnatal day 0.5 (P0.5) but sharply gear down after P7 (Figure 1A) (Shigeoka et al., 2016). By analyzing the public dataset GSE87064 composed of both bulk-tissue mRNA and assay for transposase-accessible chromatin (ATAC) sequencing data, across multiple developmental stages of the mouse retina (timepoints shown in Figure 1B), we first validated those large-scale transcriptional alterations indeed occurred in developing retinas of mice (Figure 1B). Notably, day P0 represents a temporal milestone for the occurrence of these changes in gene expressions (Figure 1B). Among these differentially expressed genes across mouse retinal development, multiple well-known promoting factors for the CNS axon regeneration reveal obvious trends of downregulation (Figure 1C-E). For instance, the Klf6 and Klf7 (Blackmore et al., 2012; Wang et al., 2018b) decreased sharply around stage P0 and remained relatively flat thereafter (Figure 1C). The Sox family factors, especially the Sox11 (Norsworthy et al., 2017), revealed similar trends in downregulation across retinal development (Figure 1D). Other identified factors, e.g. Lin28b, Ezh2, Myc and Uhrf, for promoting mature CNS axon regeneration (Wang et al., 2018a; Wang et al., 2023; (Belin et al., 2015) (Oh et al., 2018), all exhibited clear trends in downregulation across retinal development (Figure 1E). On the contrary, the factors that have been previously validated to inhibit axon regeneration, such as Klf4 and Klf9 (Apara et al., 2017; Moore et al., 2009), as well as Hdac5 (Cho et al., 2013), have been shown to be increased in expression in the RNA-seq across mouse retinal development (Figure 1C and E). Noteworthy, the expression of Ascl1 reached its peak in between E15 to P0, suggesting that Ascl1 contributed to pro-neuronal transition from progenitors with high stemness to neurons. Similar to Sox11 and Lin28, the expression of Ascl1 almost decreased to baseline after P3, suggesting strong correlation with the cessation of developmental axon projection (Figure 1F). In line with these whole retina bulk-tissue sequencing data across development, the RNA-seq (GSE185671) of the purified retinal ganglion cells (RGCs) (Shekhar et al., 2022) also confirmed that Ascl1 reached a peak expression around E16 to P0 (Figure 1G).

**Figure 1.**
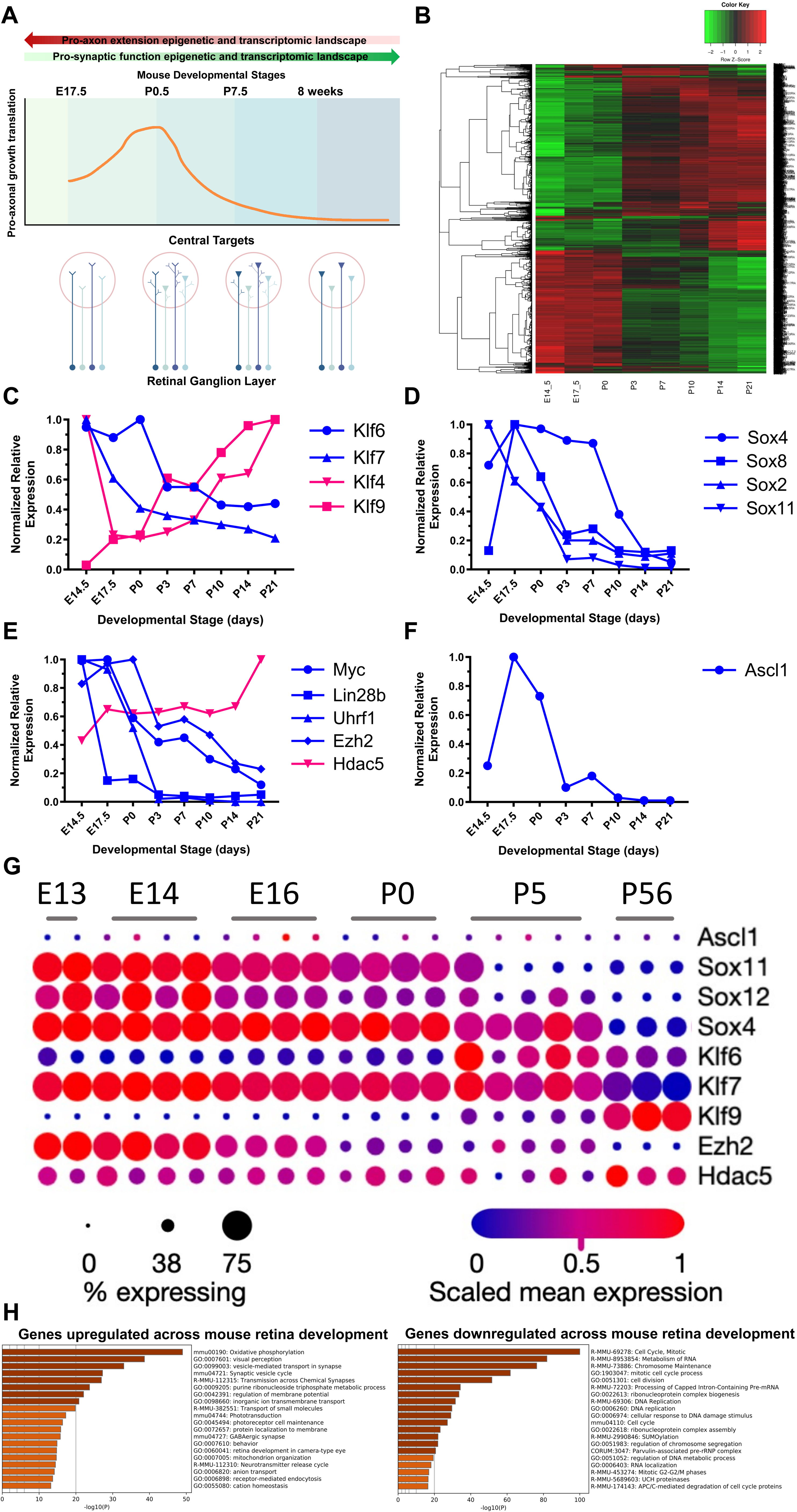
The state of retinal ganglion cells projection is in correspondence with differential gene expression across mouse retinal development. **(A)** Diagram showing the expression of axon guidance-related molecules and functional stages of RGC axon growth are closely related to the mouse embryonic and postnatal developmental stages. **(B)** Heatmap illustrating the differentially expressed genes dynamics and patterns across the mouse retinal development (public dataset GSE87064). **(C-F)** Line graphs showing the dynamics of *Ascl1* expression across retinal development is consistent with several known promoting factors to axon regeneration, while the trend of *Ascl1* expression is in opposite to the known inhibitive factors to axon regeneration. The relative mRNA expression level was normalized to the highest expression point. **(G)** Dot plot exhibiting variable gene expression and cell number coverage of representing regenerative promoting factors across developmental stages of Thy1-dependent purified RGCs (raw scRNA-seq data is from the public dataset GSE185671). **(H)** Gene ontology (G) summarizing the functions of genes upregulated (left) or downregulated (right) across mouse retina development.

The gene ontology (GO) analysis revealed numerous biological processes associated with mature neuron functions, for example the synaptic transmissions and membrane potential maintenance, were enriched by the upregulated genes across retinal development (Figure 1H, left panel). On the contrary, GO annotations associated with neurogenic proliferation, cell division and chromosome maintenance, were enriched for the downregulated genes (Figure 1H, right panel). Although the expression dynamics of Ascl1 across retinal development is in consistency with other regeneration promoting factors, it remains unclear if Ascl1 acts as a pioneering transcription factor that actually regulates regeneration associated genes (RAGs). We next performed computational analysis on the assay for transposase-accessible chromatin sequencing (ATAC-seq) dataset GSE87064 of the mouse retina development to further confidently predict that Ascl1 potentially binds to gene loci or gene enhancer regions associated with the known regeneration-associated genes (RAGs). The ATAC-seq analysis and the genome footprint-based predictions on TF-to-promoter and TF-to-enhancer interactions were performed following a workflow and algorithm previously published by our group (Gao and Qian, 2020; Gao et al., 2022; Jiang et al., 2022). We first profiled the ATAC seq dataset across retinal development and identified differentially accessible chromatin regions, for example, which changed from euchromatin (“open”) to heterochromatin (“closed”), or vice versa (Figure 2A-D). Most of these differentially accessible chromatin regions are gene promoters or intergenic regions (Figure 2B). The gene enhancers within intergenic regions were identified using the overlap areas between histone marker H3K27ac and H3K4me1 (Figure 2C) based on our method previously published (Gao and Qian, 2020; Gao et al., 2022; Jiang et al., 2022). With the gene promoters and enhancers identified, we revealed that these chromatin regions were relatively turned from open to closed during the mouse retinal development, which was consistent with the progressive loss of stemness and the establishment of terminally differentiated mature neurons and non-neuron cells. Next, we sought to predict the promoters and enhancers of RAGs were regulated by what transcription factors, using a ATAC footprint (DNA binding motif)-based method. A gene list of terminal RAGs, representing the transcriptome of high axon regenerative state, were created using the integration of some public microarray and RNA-seq datasets, which include Klf6, Klf7 or Sox11 overexpression in cortical neurons or purified retinal ganglion cells, also downregulated genes upon Klf9 overexpression (Norsworthy et al., 2017) (Galvao et al., 2018; Sun et al., 2011). The potential axon regeneration-promoting TFs were scored based on the quantity of promoters or enhancers of RAGs regulated. Specifically, Ascl1 received a score of 31 at the RAG promoters, together with some well-known TFs such as Sox11 scored 4, Fos scored 6 and Id2 scored 14. For the RAG enhancer regulation, Ascl1 received a score of 25, together with other known TFs such as Sox11 scored 1, Fos scored 5 and Id2 scored 5. It indicates our sequencing-based computational prediction can successfully identify the functionally validated TFs that promote mature CNS axon regeneration, also suggesting the predicted novel target Ascl1 highly likely act in a similar manner to do so.

**Figure 2.**
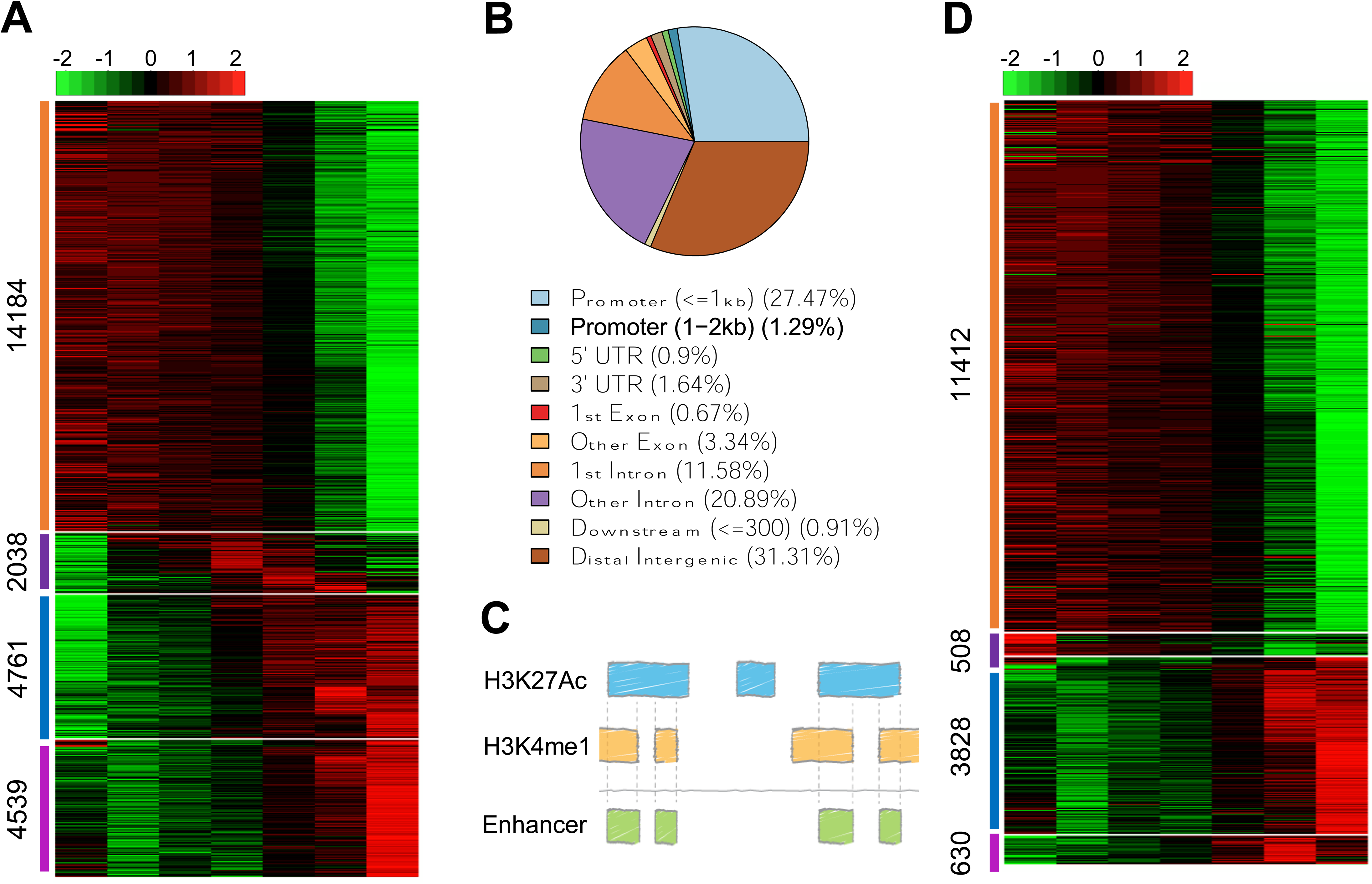
Mouse retinal development corresponds to differential chromatin accessibility. **(A)** Heatmap of ATAC-seq showing the transition of developmentally open or closed genomic regions (peaks) in mouse retina across embryonic and postnatal stages. The numbers on the left of each differential pattern indicates the quantification of peaks. **(B)** Pie chart revealing the natures of genomic regions of the differential peaks shown in (A). **(C)** Diagram illustrating the method using the ChIP-seq of H3K27Ac and H3K4me1 histone markers for characterizing active enhancer regions. **(D)** Heatmap showing the differentially accessible patterns of gene enhancer regions. The numbers on the left of each differential pattern indicates the quantification of enhancer peaks. All ATAC-seq and ChIP-seq raw data were obtained from the public dataset GSE87064.

### Ascl1 overexpression in mouse retina promotes post-injury axon regeneration and preserves retinal ganglion cell survival

To functionally investigate the effect of Ascl1, we employed the optic nerve crush (ONC) in mice, which reveals similar transcriptome with glaucoma (Wang et al., 2021; Yang et al., 2007) and is a prevailingly applied *in vivo* model to investigate the death and survival of CNS neurons (Duan et al., 2015; Guo et al., 2021; Li et al., 2022; Tran et al., 2019; Welsbie et al., 2013). The mouse retinas were transduced with AAV2-Ef1a-mAscl1-FLAG individually by intravitreal injection for overexpressing (O/E) Ascl1. Injection of AAV2-CMV-GFP was used as the control group. After 14 days of viral transduction, optic nerves across all groups were crushed and allowed 16 days for axon regeneration. The results showed that compared to the control group, Ascl1 O/E significantly promoted axon regeneration past the optic nerve crush site (Figure 3A and B), indicating Ascl1 can steer the mature RGCs into a regeneration-conducive state. Notably, Ascl1 O/E allowed for the regrowth of axons significantly distal of the crush site (Figure 3A, bottom nerve red arrows), a phenomenon cannot be observed due to the intrinsic though weak regeneration of CNS neurons. Therefore, overexpression of Ascl1 in mouse retina induces robust axon regeneration after optic nerve crush, likely through the Ascl1-mediated downstream regulation of RAGs.

**Figure 3.**
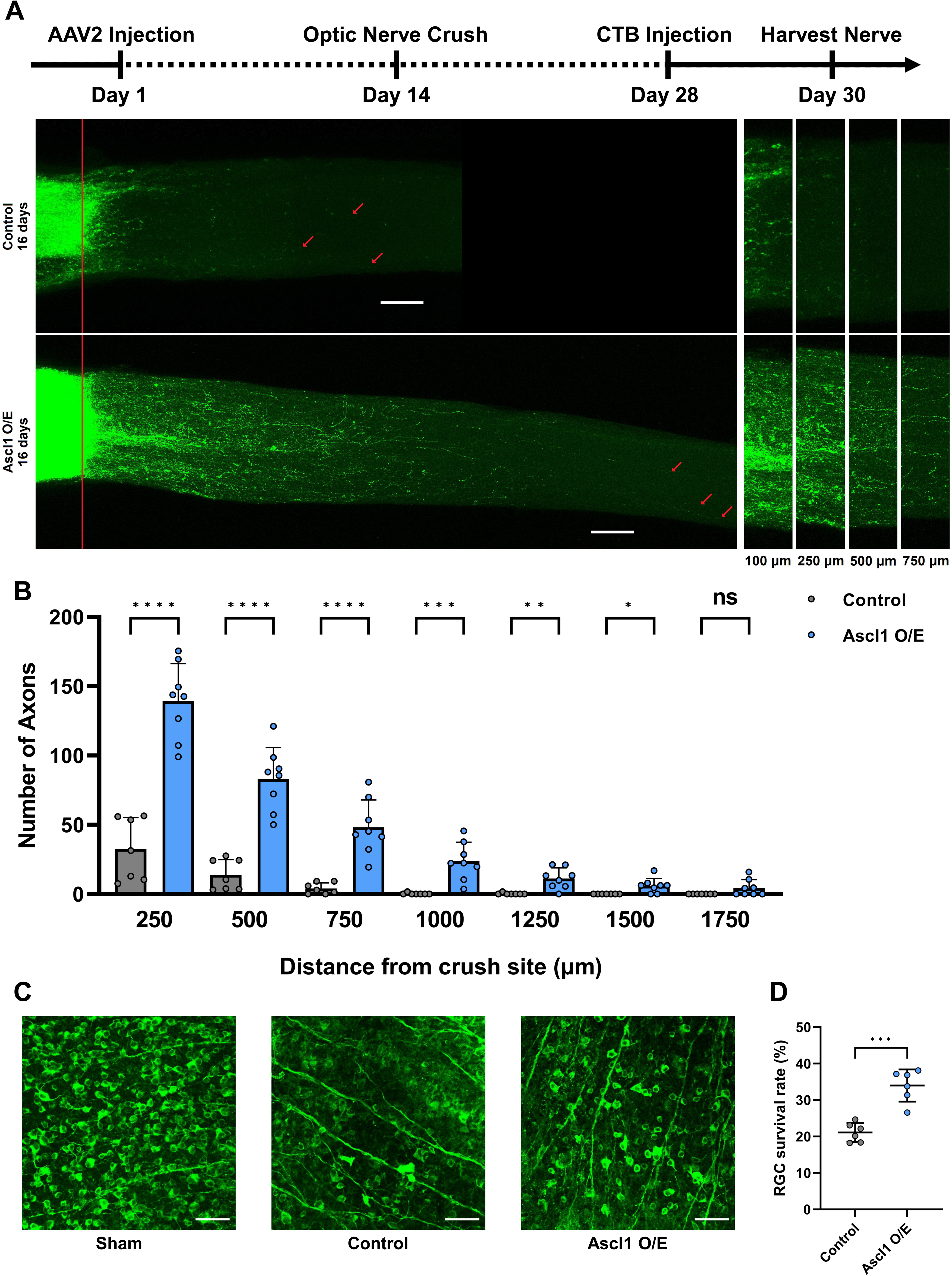
Ascl1 overexpression in adult mouse retina promotes post-injury axon regeneration and preserves RGC survival rate. **(A)** Top: timeline of the experiment. Middle: representative images showing that Ascl1 O/E in adult mouse retina promotes substantial axon regeneration in optic nerves 16 days after optic nerve crush compared to the control group. The red line indicates the crush sites. The red arrows indicate the longest axons of each nerve. Scale bar is 100μm in length. Bottom: Enlarged images of sections of nerves at 100 μm, 250 μm, 500 μm, and 750 μm distal of the crush sites. **(B)** Bar chart showing quantification of (A). A dot represents an individual optic nerve [n = 7 mice in the 16 day injection-to-crush control group, n = 8 mice in the 16 day injection-to-crush *Ascl1* O/E group]. Unpaired *t*-tests were applied at all distances [*p* < 0.0001 at 250 μm, 500 μm, and 750 μm; *p* = 0.0005 at 1000 μm; *p* = 0.002 at 1250 μm; *p* = 0.0104 at 1500 μm; *p* = 0.0727 at 1750 μm]. **(C)** Representative image fields of whole mount retinas showing that Ascl1 O/E in adult RGCs significantly increased RGC survival rate 16 days after optic nerve crush compared to the control group. From left, sham no injury retina, control group retina 16 days after optic nerve crush, Ascl1 O/E retina 16 days after optic nerve crush. Whole-mount retinas were stained with anti-βIII tubulin (Tuj1 antibody, in green fluorescence). White scale bar is 50μm in length. **(D)** Scatter plot showing quantification of (C). An unpaired *t-*test was applied [*p* = 0.0001]. n = 6 mice in the 16 day injection-to-crush control group, n = 6 mice in the 16 day injection-to-crush *Ascl1* O/E group). ns not significant. * *p* < 0.05, \*\**p* < 0.01, *** *p* < 0.001, **** *p* < 0.0001.

Furthermore, neuronal cell death remains a critical obstacle to functional recovery following CNS injury. In the adult mammalian retina, around only 20% of RGCs survive 14 days after optic nerve crush (Wang et al., 2018a). In this work, we showed that Ascl1 O/E significantly increased RGC survival rate 16 days after the optic nerve crush injury (Figure 3C and D), implying the role of Ascl1 as a neuroprotective factor. Experimentally, the neuroprotective effect of Ascl1 has not been fully exhibited due to the limit in cell coverage of the viral transduction (Figure 4).

**Figure 4.**
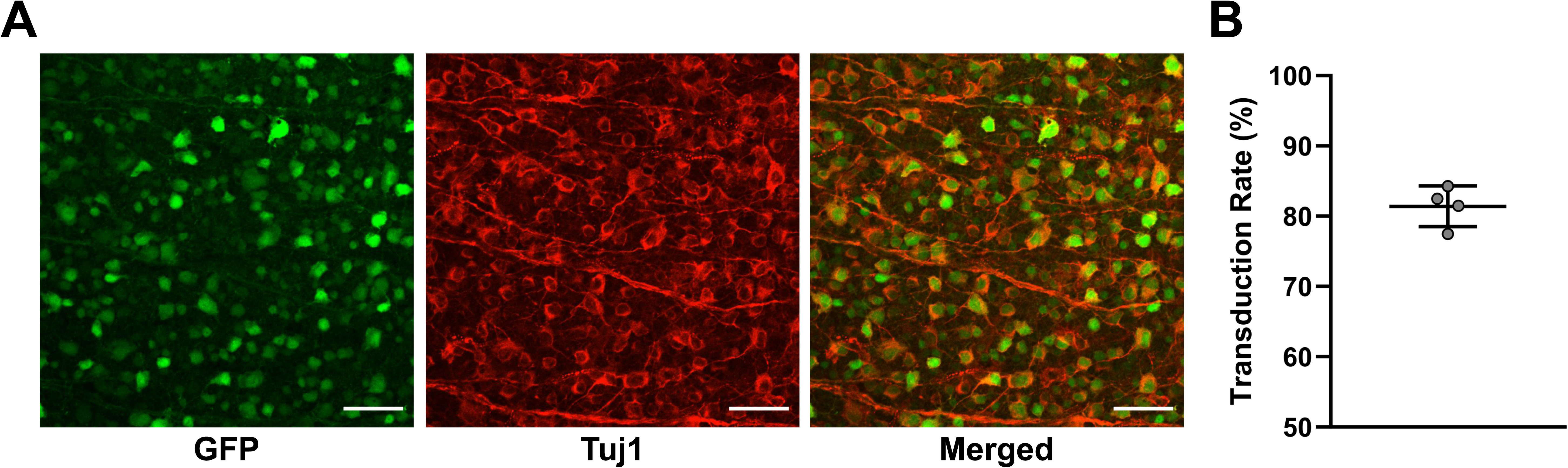
The quantification of AAV2 virus transduction rate in RGCs. **(A)** Representative image field of a whole mount retina showing AAV2 GFP transduction rate in RGCs. From left, GFP single channel, Tuj1 single channel, combined Tuj1 (red) + GFP (green) channels. Whole-mount retinas were stained with anti-βIII tubulin (Tuj1 antibody, in red channel) antibodies. White scale bar is 50μm in length. **(B)** Quantification of the transduction rate of AAV2-GFP in RGCs in (A). The average transduction rate was 81.416 ± 2.496%. The bars on the graph represent standard deviation. A dot represents an individual retina [n = 4 mice].

### Ascl1 overexpression selectively upregulates a known axon regenerative factor but not others

In order to explicate the molecular mechanisms of Ascl1-mediated RGC axon regeneration, mouse retinas were transduced with AAV2-Ef1a-Ascl1-FLAG. Retinas were harvested 14 days after viral transduction, followed by the cryosection of the retinal tissues. Samples were then immuno-stained against multiple well-known mediators of CNS axon regeneration (Sox11, Sox9, Stat3, Klf4, and phosphor-GSK3β) to see if any of these cell signals were mediated by Ascl1 O/E. We found that only the Sox11 fluorescent intensity in the retinal ganglion layer (GCL) significantly upregulated compared to the control group (Figure 5A), while other known regeneration-associated cell signals were not significantly changed by Ascl1 O/E in mouse retina (Figure 5B-E). The axon regeneration favorable Stat3 was not increased, nor the regeneration inhibitory Klf4 or phosphorylated GSK3β was downregulated. These results indicate that Sox11 may be the downstream of Ascl1 signaling and thus mediates Ascl1’s pro-regenerative effects.

**Figure 5.**
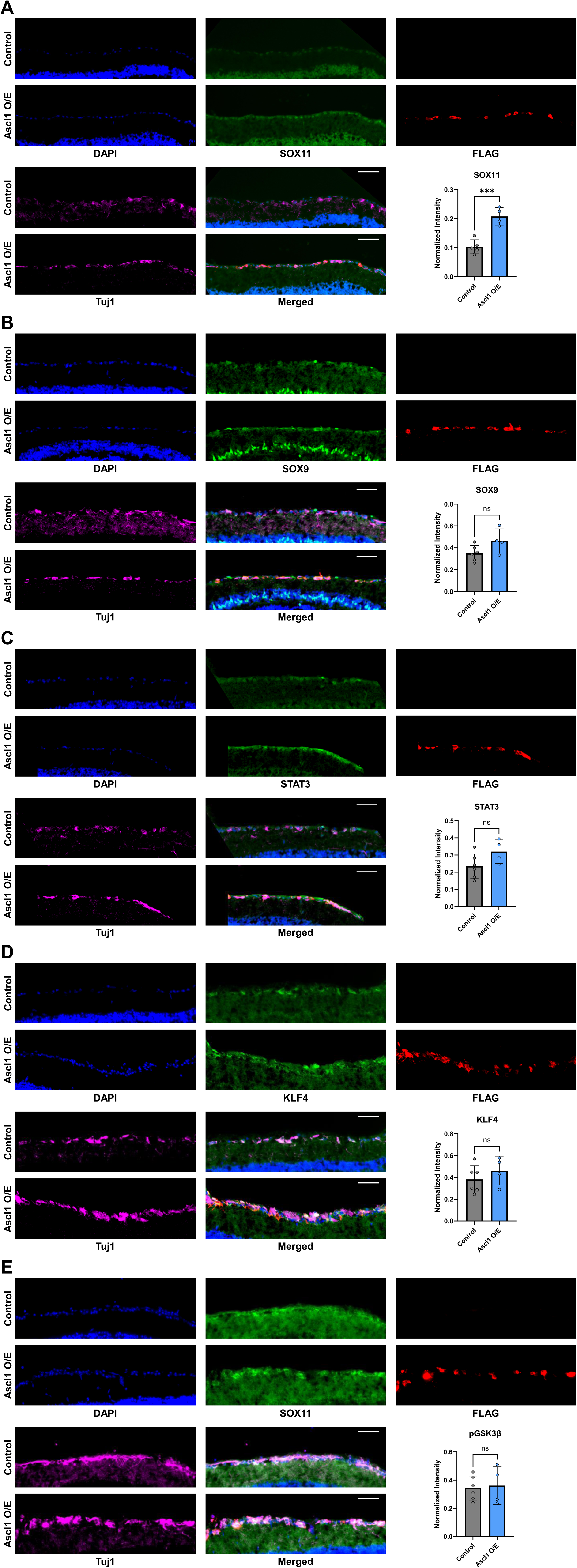
Ascl1 overexpression upregulates Sox11 but not other axon regeneration associated factors in adult mouse RGCs. **(A)** Representative images indicating greater Sox11 immuno-staining intensity in the RGC layer under Ascl1 O/E two days after ONC. Retinal cryosections were stained with DAPI (blue), anti-SOX11 (green), anti-FLAG (red) and anti-tubulin β3 (far-red) antibodies. The white scale bar is 50 μm. Bottom right is quantification of normalized intensity (unpaired T-test, *p =* 0.0007, n = 5 retinas for control, n = 4 retinas for Ascl1 O/E). **(B)** Representative images indicating no significant difference in Sox9 intensity between Tuj1+ and FLAG+ Ascl1 O/E cells and Tuj1+ cells (unpaired T-test, *p =* 0.0825, n=6 retinas for control, n=4 retinas for Ascl1 O/E). **(C)** Representative images indicating no significant difference in Stat3 intensity between Tuj1+ and FLAG+ Ascl1 O/E cells and Tuj1+ control cells (unpaired T-test, *p =* 0.0990, n=6 retinas for control, n=4 retinas for Ascl1 O/E). **(D)** Representative images indicating no significant difference in Klf4 intensity between Tuj1+ and FLAG+ Ascl1 O/E cells and Tuj1+ control cells (unpaired T-test, *p =* 0.3766, n=6 retinas for control, n=4 retinas for Ascl1 O/E). **(E)** Representative images indicating no significant difference in phospho-GSK3β intensity between Tuj1+ and FLAG+ Ascl1 O/E cells and Tuj1+ control cells (unpaired T-test, *p =* 0.7996, n=6 retinas for control, n=4 retinas for Ascl1 O/E). ns not significant. *** *p* < 0.001. Representative images from the Ascl1 O/E were normalized such that *I*_*background*_ was equal to that of the control condition.

## DISCUSSION

After development into maturity, CNS neurons transition from an inducible growing condition to a permanent regenerative incapability (Shewan et al., 1995). With the discovery and achievement of artificially inducing terminally differentiated somatic cells backward to the inducible pluripotent stem cells (iPSCs), a consensus has been reached that the cellular identities and states, for instance the ability in axon regeneration, may be convertible bidirectionally (Gaub et al., 2010; Lindner et al., 2013; Wahane et al., 2019). In this study, the post-injury axon regeneration in the adult mouse retinal ganglion cells with Ascl1 overexpression provided additional evidence that the regulators of epigenetic reprogramming, for example the pioneer transcription factor Ascl1, had the potential to drive a large-scale epigenetic and transcriptional transition which enabled mature CNS neurons to regenerate injured axons.

The retinal cryosection immunostaining data implicate Sox11 as one of the RAGs downstream of Ascl1. Sox11, also with pioneering properties, is a member of the SRY-related high mobility group box transcription factors and has been implicated as a key mediator in neural stem cell proliferation, neurogenesis, and migration (Bergsland et al., 2006; Wang et al., 2013). Importantly, endogenous Sox11 signaling is necessary for the spontaneous neuro-regenerative process in mature peripheral nervous system following injury (Jankowski et al., 2009), while the forced Sox11 overexpression has been shown to promote axon regeneration in mature saphenous nerve (Jing et al., 2012), optic nerve (Norsworthy et al., 2017), and the corticospinal tract (CST) mouse models (Wang et al., 2015). Related to the previously identified cell signaling, Sox11 is downstream of Ascl1 and other bHLH family TFs such as Ngn2 and Math3 during the neurodevelopment (Bergsland et al., 2006). Furthermore, Ascl1-mediated astrocyte-to-neuron conversion was accompanied with an increase in the expression of Sox11 (Rao et al., 2021). In line with the mechanistic exploration in this work, all these previous studies suggest that Ascl1 may be the upstream of the SOX11 pathway.

In the paradigm of axon regeneration, it is known that many RAGs are either proto-oncogenes, or participate in signaling pathways initially discovered in tumorous cells, which may include but not limit to the abnormal expression of p300, HDACs, PTEN, and Klf family TFs found in a myriad of different types of cancers (Liu et al., 2019; Milella et al., 2015; Tetreault et al., 2013). Also, the misfunction of PI3K/Akt signaling is implicated in a variety of cancers (Janku et al., 2018). The prediction of Ascl1 is based on the computational analysis of neuron-specific developmental and maturation processes, suggesting that Ascl1 is potentially an innate regulator preset for the nervous system. Additionally, with regards to the epigenetic reprogramming for achieving mature CNS axon regeneration, the pTFs such as Ascl1 possess great advantages over the global alterations induced by DNA motif independent epigenetic modifications (e.g. the HATs and HDACs). The pTFs have been thought to specifically bind to DNA sequences on the chromatin (Zaret and Carroll, 2011), while many HATs and HDACs change the chromatin structure across the entire genome (Katan-Khaykovich and Struhl, 2002), thus potentially altering the expression and accessibilities of many genome loci that actually counteract or do not participate in axon regeneration, leading to reduced net potency for axon regeneration or strong off-target unintended effects.

## MATERIALS AND METHODS

### Mice

All animal experiments were conducted in accordance with the protocol approved by the Institutional Animal Care and Use Committee of Johns Hopkins University. 6-8 week old female CD-1 mice and male C57BL/6J mice were used in animal experiments. The CD-1 wild type mice were ordered from Charles River and the male C57BL/6J mice were ordered from the Jackson Laboratory. All animal surgeries were performed under anesthesia induced by intraperitoneal injection of avertin (20mg/mL) diluted in sterile saline. Details of the surgeries are described below.

### Optic nerve regeneration model

Cavities were made just posterior of the ciliary bodies in the right eyes of mice by inserting a 28-gauge needle at a 45-degree angle into the sclera half the length of the needle bevel. 1.5 μL of adenosine-associated virus 2- (AAV2) packaged transgene was injected through the cavity into the vitreous humors using a Hamilton syringe (Model 75 RN Syringe, 30-gauge needle). Care was taken to prevent lens, scleral, and retinal damage by either needle or syringe insertion. 14 days after viral injection, the right optic nerves were exposed laterally by retracting the retractor bulbi muscle and were crushed with Dumont #5 fine forceps (Fine Science Tools) for 10 s at approximately 1.5 mm behind the optic disc. The crush intensity was controlled by applying just enough pressure to ensure the two tips of the forceps were fully closed. 14 days after the optic nerve crush, 1.5 μL of cholera toxin subunit B (CTB) conjugated with Alexa Fluor 594 (2 mg/ml, Thermo Fisher Scientific) was injected into the right vitreous humors of the same mice (same procedure as viral injection) to allow for anterograde tracing of RGC axons in the optic nerves. 2 days after CTB injection, the mice were sacrificed by transcardial perfusion. The right eyeballs were dissected out with the portion of the optic nerve proximal of the optic chiasm remaining attached. The eyes with optic nerves attached were post-fixed in 4% paraformaldehyde (PFA) overnight at 4°C. The optic nerves were then carefully dissected out in 1x phosphate buffered saline (PBS) to ensure the end of the optic nerve most proximal to the optic disc included the crush site. Optic nerves were then washed three times with PBS to flush out residual PFA. AAV2-GFP was purchased from SignaGen Laboratories. AAV2-EF1a-Ascl1-Flag was packaged by Viagene. All viruses used had titers of 0.5 × 10^13 constructs/mL.

### Optic nerve dehydration and clearing

Fixed optic nerves were incubated in stepwise concentrations of tetrahydrofuran (TFH) (steps: 50%, 70%, 80%, 100%, and 100%, v/v % in distilled water, Sigma-Aldrich) in a twelve-well glass plate to dehydrate the nerves. For the final step, nerves were incubated in benzyl alcohol/benzyl benzoate (BABB, 1:2 in volume, Sigma-Aldrich) for tissue clearing. All incubations were performed on an orbital shaker at room temperature for 20 minutes each step, with six steps in total. The dehydrated and cleared optic nerves were mounted in BABB in specialized silica gel wells. The plate was covered in aluminum foil through the entire dehydration and clearing to reduce photobleaching of the optic nerves.

### Imaging of RGC axon regeneration

Mounted whole-mount nerves were imaged with a LSM 800 confocal microscope (Zeiss, 20x air objective) with Zeiss ZEN Blue software. The z coordinates of each nerve were first defined by manually adjusting the imaging plane upwards or downwards until signal was no longer observed, demarcating the upper and lower z coordinates, respectively. Then, the Z stack function was used to acquire the range of the z coordinates of the nerve through slices (2 μm) to produce a stack spanning from the lower to upper z coordinates. The x-y coordinates of each nerve were defined by manually tracing the nerve to create imaging tiles that fully encompassed the x-y plane of the nerve. The tiling function (10% overlap) was applied between adjacent tiles to image the whole nerve, such that tiles of each 2 μm-thick slice were stitched to create a complete slice image of the entire nerve. Complete slices could then be combined along the z-axis using Z-stack function.

### Analysis of RGC axon regeneration

Quantification of regenerating axons in the optic nerve was performed as previously described (Wang et al., 2018a). Starting from the lower z coordinate, 10 continuous slices were merged along the z-axis to generate a series of 20 μm-thick sections across the entire z-range of the optic nerve. Starting from the crush site (0 μm), distances at 250 μm intervals were drawn, up to 1750 μm from the crush site. At each distance for each 20 μm-thick sections, the number of CTB-labeled axons was counted. For each nerve, the total number of axons at each distance from the crush site was obtained by summing together the number of axons of all 20 μm-thick sections. The maximum diameter of the nerve at each distance was taken to calculate number of axons per micrometer of nerve width, then averaged over all sections. Sad, the total number of axons at distance d in a nerve with a maximum radius of r, was estimated by summing over all optical sections with a thickness of t (20 μm): Sad = πr^2 x (average axons/mm)/t.

### Immunohistochemistry of whole-mount retinas

After overnight post-fixation in 4% PFA at 4°C, eyeballs were dissected from the attached optic nerves. Retinas were then carefully dissected from the eyeballs 1x PBS and washed three times in 1x PBS to flush out residual PFA. Retinas were then cut into petal shape (4 opposing radial incisions) and blocked in blocking buffer (1% Triton X-100, 10% goat serum in 1x PBS) for 1 hr at room temperature. Retinas were then stained with primary antibodies (1:500 mouse anti-Tubulin β3, BioLegend) at 4°C, followed by the corresponding Alexa Fluor-conjugated secondary antibodies (1:500, Thermo Fisher Scientific) for 1 hr at room temperature. All antibodies were diluted in blocking buffer. Following both primary and secondary antibody incubation, the retinas were washed in washing buffer (Triton X-100 0.3% in 1x PBS) for four times, for 15 min each time. After the last wash, the retinas were mounted onto slides with ProLong Gold Antifade Mountant with DAPI (Invitrogen).

### Analysis of RGC Survival Rate

Eight image fields were selected randomly and imaged from the peripheral regions of each whole-mount retina with a LSM 800 confocal microscope (Zeiss, 20x air objective). RGC survival rate was calculated by dividing the average number of Tuj1+ cells from all eight fields in the injured retinas by that in the uninjured retina. Injured retinas (control and *Ascl1* O/E) were performed under the same timeframe as with the optic nerve regeneration model (16 days after optic nerve crush).

### Analysis of RGC Transduction Rate

Uninjured retinas were dissected 14 days after AAV2-GFP injection and fixed in whole-mount, as described above. Six image fields were selected randomly and imaged from the peripheral regions of each whole-mount retina with a LSM 800 confocal microscope (Zeiss, 20x air objective). RGC transduction rate for each image was calculated by counting the number of Tuj1 positive + GFP positive cells and dividing by the number of Tuj1 positive cells.

### Immunohistochemistry of retinal cryosections

Retinas were dissected and washed in PBS as described above. Washed retinas were cryoprotected in stepwise sucrose incubations (steps: 10%, 20%, 30%, w/v % in 1x PBS). All steps were performed on the orbital shaker at room temperature until the retinas sunk. Cryoprotected retinas were transferred to plastic molds filled with Tissue-Tek O.C.T. Compound (Sakura), flash frozen on dry ice, and stored at -80°C until cryosectioning. Retinas were then cryosectioned laterally at 10 μm section thickness onto Fisherbrand Superfrost Plus microscope slides (Thermo Fisher) and stored at -20°C until immunostaining. Prior to blocking, slides were warmed at 37°C for 1 hr on a slide blocker. Rectangular barriers around all the sections on a slide were created using a hydrophobic PAP pen (Newcomer Supply) during warming. Slides were then blocked in blocking buffer (0.3% Triton X-100, 10% goat serum in 1x PBS) for 1 hr, stained with primary antibodies overnight (1:500 mouse anti-Tubulin β3, BioLegend; 1:500 rabbit anti-Sox11, abcam; 1:500 rabbit anti-Sox9, Sigma-Aldrich; 1:250 rabbit anti-Stat3, CST; 1:500 rabbit anti-KLF4, Millipore; 1:250 rabbit anti-pGSK-3β (Ser9), CST; 1:500 rat anti-FLAG, Novus) at 4°C, followed by the corresponding Alexa Fluor-conjugated secondary antibodies (1:500, Thermo Fisher Scientific) for 1 hr at room temperature. All antibodies were diluted in blocking buffer. Following both primary and secondary antibody incubation, the retinas were washed in washing buffer (Triton X-100, 0.3% in 1x PBS) for four times, for 15 min each time. After the last wash, the slides were sealed with ProLong Gold Antifade Mountant with DAPI (Invitrogen).

### Analysis of retinal cryosections

Fluorescent images of the retina cryosections were acquired with an Axiovert 200M inverted fluorescent microscope (Zeiss, 20x air objective). For the Ascl1 O/E condition, regions of interest (ROIs) in ImageJ were drawn around Tuj1+ and FLAG+ cells that were counterstained with DAPI. For the control condition, ROIs were drawn around Tuj1+ cells that were counterstained with DAPI. Fluorescent intensity for an image was measured as the mean pixel intensity in ROIs, weighted for the area of ROIs. Average fluorescent intensity was calculated as from 6 images per retina. Normalized intensity was calculated as follows *I*_*norm*_ = (*I*_*average*_ − *I*_*background*_)/*I*_*max*_.

### Quantification and Statistical Analysis

Statistical analyses were completed with GraphPad Prism 9 and the significance level was set as *p* < 0.05. Data are represented as mean ± standard deviation (error bars) unless specifically stated. For comparisons between two groups, two-tailed unpaired t tests were used. All details regarding statistical analyses, including the tests used, *p-*values, exact values of n, definitions of n, are described in figure legends.

**Table 1.**
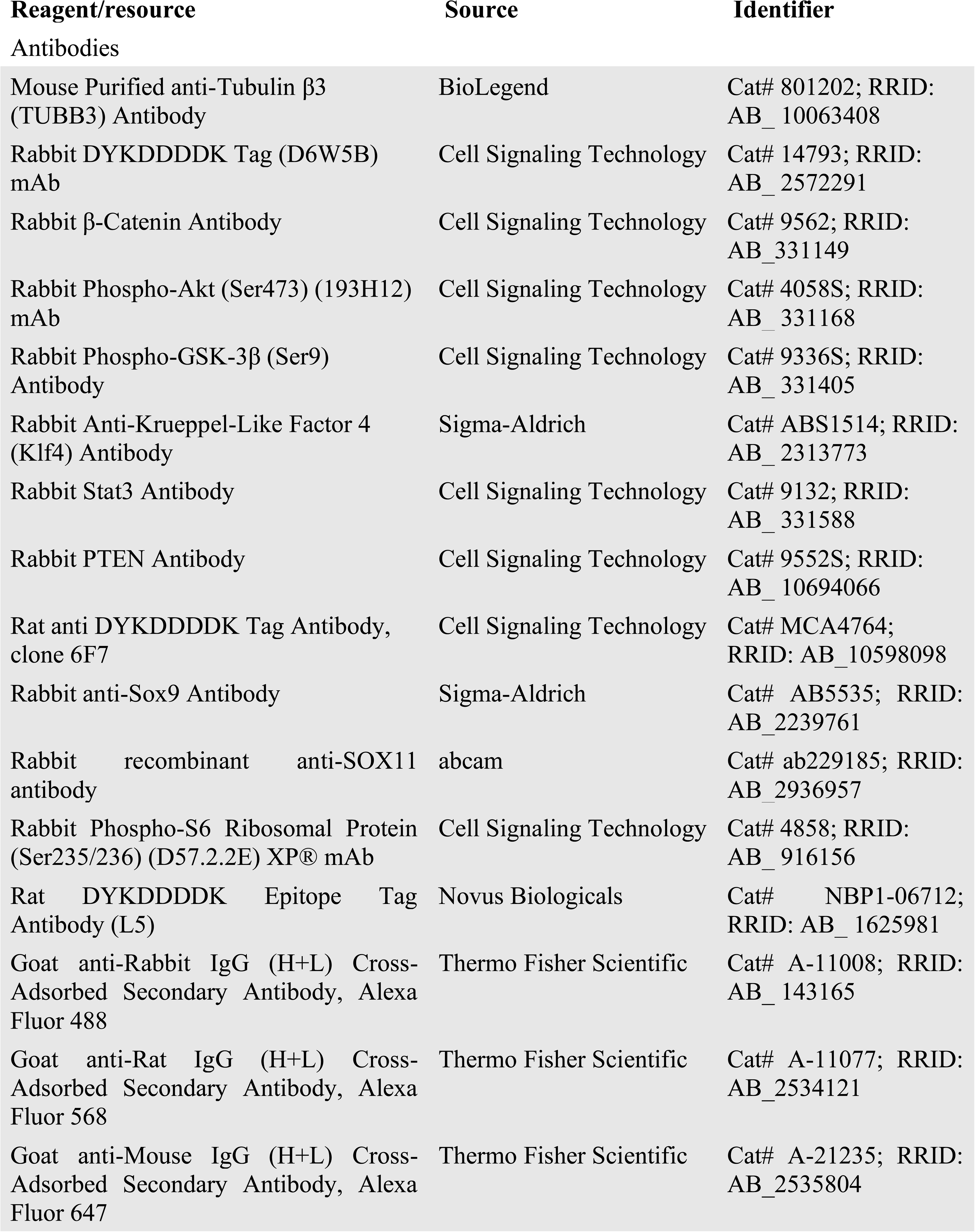
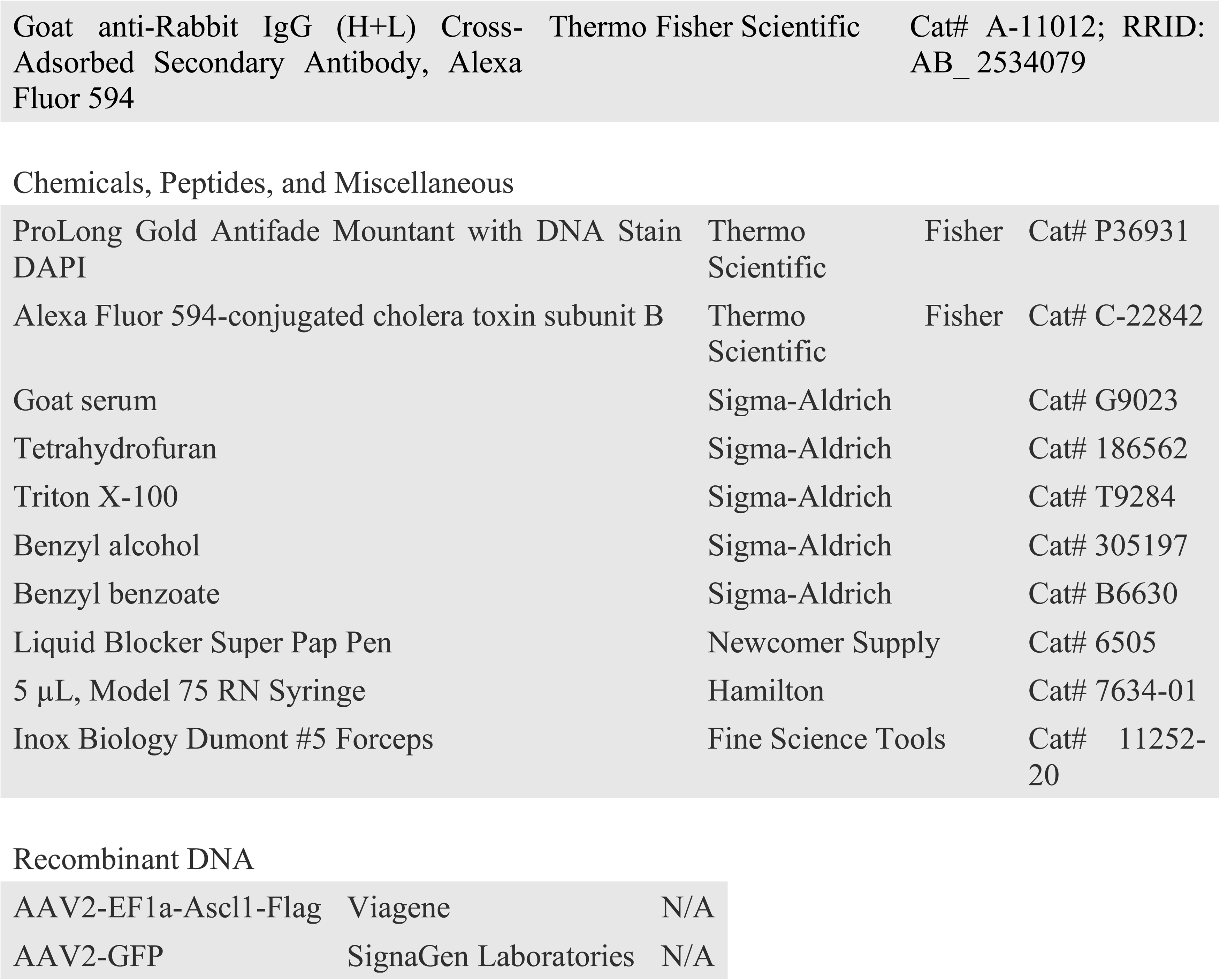
Key Resources.

